# Development of Int-Plex@ binary memory switch system: plant genome modulation driven by large serine-integrases

**DOI:** 10.1101/2024.01.11.575089

**Authors:** Marco Antônio de Oliveira, Rayane Nunes Lima, Lilian Hasegawa Florentino, Mariana Martins Severo de Almeida, Fernando Lucas Melo, Raquel Vasquez Bonnet, Cristiano Lacorte, Elibio Rech

## Abstract

The comprehension of virus-host interactions has allowed numerous advances in developing biotechnological methodologies for plant genome editions, constituting a promising path for plant genetic engineering. Among these advancements, phage-encoded large serine-integrases have emerged as noteworthy tools to modulate plant metabolic pathways by inserting, excising, or inverting DNA stretches in a reversible and specific way. The present work shows the foundation of the Int-Plex@ (INTegrase PLant EXpression) binary memory switch system, which consists of the application of four distinct orthogonal prophage large serine-integrases (Int) (BxB1, phiC31, Int13, and Int9) as an input trigger mechanism for the inversion or excision of genomic DNA. The memory genetic switch is divided into the excision module and the inversion module. The excision module is activated by BxB1 or phiC31 enzymes (input). In this case, the DNA sequence flanked by its attachment sites is excised from the genome (output). The inversion module is activated by Int9 or Int13 (input). The inverted *mgf* gene sequence is flipped to its functional coding sequence, and the switch output is mGFP. Moreover, prokaryotic-based cell-free *in vitro* transcription-translation reactions (TxTl) were used as a fast platform for testing Ints *attB/P in tandem* site activity. Furthermore different plasmid delivery strategies for plant cell Int heterologous expression were tested: leaf tissue agroinfiltration of *Agrobacterium tumefaciens* transformed with binary plasmids and a biolistic system. After each treatment, the edited genomic DNA sequences were amplified and verified by Sanger and Nanopore sequencing. Despite the challenges of using Ints, the potential benefits are significant and deserve deeper exploration and development. The Int-Plex@ binary genome memory switch system can be applied to produce genetic circuits combined with omics tools and sgRNAs to engineer and modulate plant metabolic pathways temporally and reversibly.

## 1. INTRODUCTION

Over the last several decades, major advances in plant synthetic biology have emerged from significant discoveries based on the understanding of the co-evolutionary arms race between viruses and their hosts. These ongoing battles have led to the evolution of intricate and complex molecular interactions that were exploited to create novel biotechnological tools based on the bottom-up approach for plant genome modulation. Examples include (I) bacteria restriction endonucleases and phage DNA ligases, which allowed the revolution of recombinant DNA technology and are used in BioBrick^TM^ assembly and modern one-step DNA assembly techniques^1–7^; (II) viral vectors for gene delivery^8,9^; (III) virus-induced gene silencing (VIGS)^10,11^; (IV) the RNA interference (RNAi) technology, which is widely used in gene down-regulation studies^12–14^; and (V) CRISPR-Cas systems, which has recently been widely applied in crop genome editing technology^12,15–18^. However, these last two technologies are irreversible and permanent; once the plant gene is silenced or edited, it cannot return to its original state. Therefore, to overcome these limitations, a promising approach, originating from bacteriophage-prokaryotic interactions, is the application of filamentous phage-encoded large serine integrases (Ints), which are multifunctional recombinases with the potential to edit DNA sequences and also serve as efficient *in vitro* DNA assemblers^19–22^. Therefore, considering the development of a dynamic and reversible genome modulation system in plant cells, it is possible to include the orthogonal Ints in modular synthetic genetic devices based on Boolean logic gates^23–29^. Ints catalyze directly and unidirectionally a site-specific insertion, deletion or 180° inversion of DNA sequence flanked by the two attachment sites (*att*), *attB* (bacteria) and *attP* (phage) depending on the design of their orientation relative to each other, which results in the formation of two new post-recombination attachment sites, *attL* (left) and *attR* (right)^19,20,25–28,30^. BxB1 and phiC31 Ints have been successfully used in plant engineering and gene stacking^29,31–34^.

Furthermore, using a bioinformatics approach, Yang *et al*. prospected 11 new Ints with their respective functional *att* sites^20^. Subsequently, Gomide *et al*. showed that those Ints function in eukaryotic cells and ranked Int9 and Int13 among them as the most effective Ints for plasmid flipping in the plant cell context^25^. Thus, the IntPlex@ binary memory switch takes BxB1, phiC31, Int9, and Int13 as input signals to control an output (genome edition) in a user-defined manner (site-specific excision or inversion). Lastly, the possibility of using distinct plasmid delivery as a general method for integrase expression was examined.

## 2. RESULTS

### 2.1 Int-Plex@ binary memory switch system design

We have assembled a multifunctional genetic switch system based on *att* site pairs from 4 Ints, namely BxB1, phiC31, Int9, and Int13. The cassette constructs consist of the reverse complement sequence of green fluorescent protein reporter gene *gfp* (*mgfp* or *degfp*) flanked by in tandem *attB* or *attP* sites from the Ints selected (Figure 1A). Promoter and terminator sequences to complete the transcription unit varied according to the *in vitro* or *in vivo* experimental model. Effector Ints were delivered individually in a separate plasmid under transcriptional control of a constitutive promoter^25,35^. This configuration allows the formation of a binary system with an Excision Module controlled by BxB1 or phiC31 recombinases designed as a NOR logic gate, in which the presence of either Int (input) prevents reporter expression due to its deletion (output) and an Inversion Module controlled by Int9 or Int13 with the structure of an XOR gate (Figure 1B). To achieve varying outcomes in DNA rearrangement depending on the Int acting on the system, we inserted the *att* sites in the construction either in the same or opposing orientation relative to their respective site counterpart. While for BxB1 and phiC31, *attB* and *attP* sites are in sense orientation, with their recombination resulting in excision of the DNA sequence flanked by them from the construction (Figure 1, C and D), Int9 and Int13 *att* sites are in reverse complement orientation, which means that upon recombination, the target DNA sequence will be 180° flipped (output), modulating the *gfp* expression (Figure 1, E and F). Although both Int9 or Int13 are capable of unidirectionally flipping the reporter gene coding sequence and activating its transcription, an essential feature of this type of logic gate is the gene set/reset, that upon concurrent or sequential introduction of both enzymes, a second inversion event will result in the return of the target DNA orientation to the original OFF state, silencing reporter expression (Figure 1, G and H). In this double-editing event, one Int will recognize its *attB/P* sites and catalyze the inversion of the reporter from its OFF state to ON state, forming the intermediate plasmid with *attL/R* sites which results in the GFP production. Instantaneously, this intermediate plasmid is the substrate for the action of the second Int, which, upon recombination of its *attB/P* sites, will result in the formation of the final plasmid with both *attL/R* sites for Int9 and Int13 with consequent recovery from the ON state to initial OFF state, silencing *gfp* expression (Figure 1, G and H).

**Figure 1.**
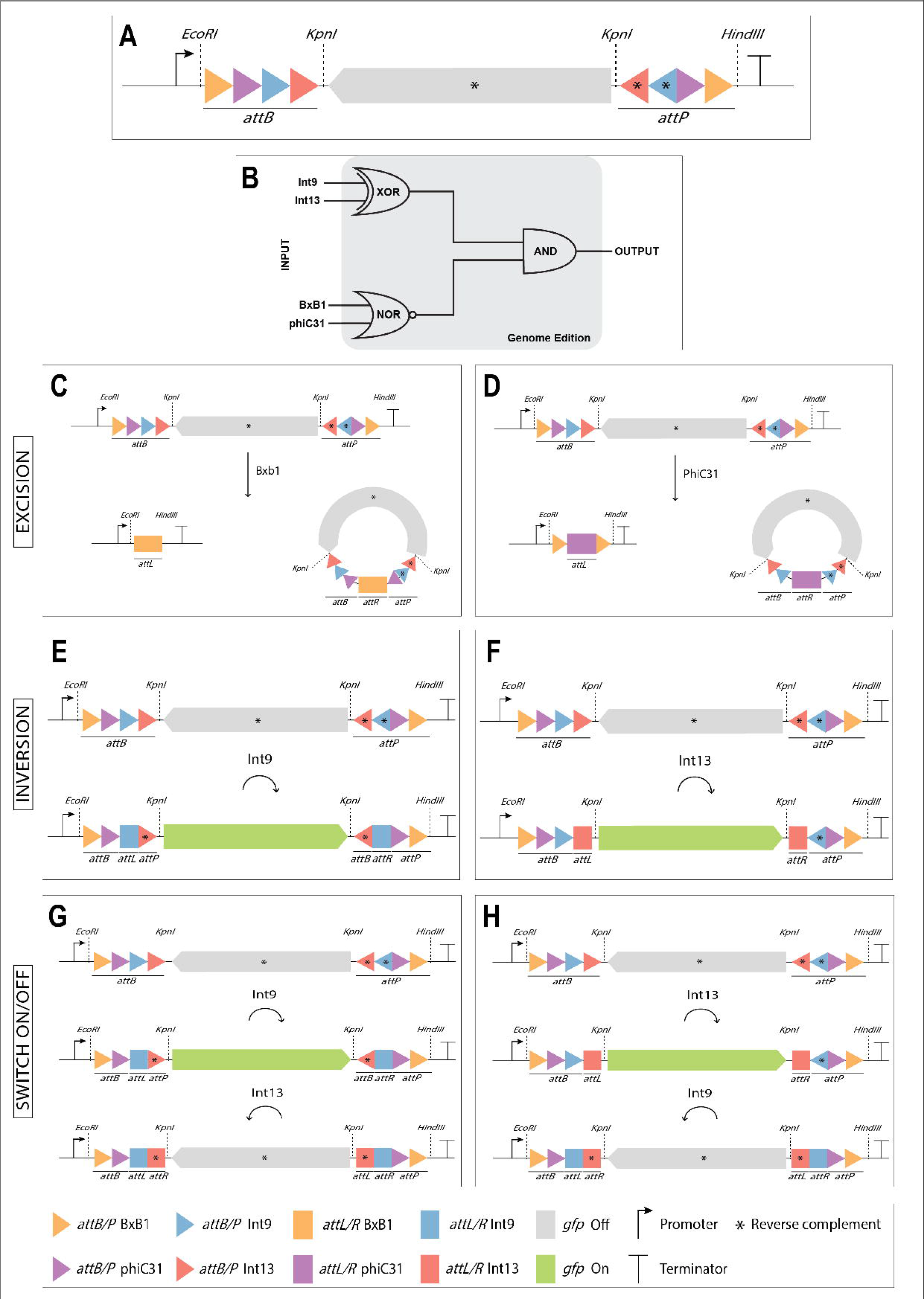
Schematic overview of the IntPlex@ memory genetic switch architecture and Boolean-based logic gates. (A) The diagram describes the architecture of the bimodular genetic switch, which consists of a *gfp* sequence in reverse orientation (gray) surrounded by four *attB* and *attP* sites (triangles) of four serine integrases: BxB1 (yellow), phiC31 (lilac), Int9 (blue) and Int13 (red). For in vitro assays, the switch contains the *degfp* gene and the switch integrated into the plant genome contains the *mgfp* gene. (B) Int9/Int13-based XOR logic gate for *gfp* orientation flipping (turn *gfp* on). Int9/Int 13 based OR logic gate for turning *gfp* on and off. BxB1/phiC31-based NOR logic gate for *gfp* excision. In the presence of Int9 or Int13, the *gfp* is inverted to its coding sequence (ON state). Moreover, in the presence of Int9 and Int 13, the *gfp* is modulated and oscillates between the on and off states. In the presence of BxB1 or phiC31 input, the *gfp* is excised. The system output is edited DNA: plasmid or plant genome. (C) The diagram shows IntPlex@ memory genetic switch operation driven by BxB1. The resulting edited DNA with the BxB1 *attL* scar (left) and the excised circular DNA molecule with BxB1 *attR* (right). (D) The diagram shows IntPlex@ memory genetic switch operation driven by phiC31. The resulting edited DNA with the phiC31 *attL* scar (left) and the excised circular DNA molecule with phiC31 *attR* (right). (E) The diagram shows IntPlex@ memory genetic switch operation driven by Int9. The sequence flanked by the Int9 *attB* and *attP* sites and, consequently, the inversion of the *gfp* to its coding sequence (GFP output). The Int13 *att* sites are also inverted and oriented towards their reverse complement. (F) The diagram shows IntPlex@ memory genetic switch operation driven by Int13. The *gfp* flanked by the Int13 *attB* and *attP* sites is flipped to its coding sequence (GFP output). (G and H) The diagram shows IntPlex@ memory genetic switch operation driven by Int9 and Int13. The *gfp* is flipped in two steps according to which integrase is added first. (G) In the first step, Int9 recognizes its *attB/P* sites and catalyzes the inversion of *gfp* from its OFF state to ON state, resulting in the formation of the intermediate sequence with *attL*/Int9 and *attR*/Int9 sites and the output GFP. Instantaneously, this intermediate sequence is the substrate for the action of Int13 that recognizes its *attB/P* sites and catalyzes the inversion of *gfp* from its ON state to OFF state, resulting in the formation of the final plasmid with both *attL/R* Int9 and *attL/R* Int13 sites. (H) In the first step, Int13 recognizes its *attB/P* sites and catalyzes the inversion of *gfp* from its OFF state to ON state, resulting in the formation of the intermediate sequence with *attL*/Int13 and *attR*/Int13 sites and the output GFP. Instantaneously, this intermediate sequence is the substrate for the action of Int9 that recognizes its *attB/P* sites and catalyzes the inversion of *gfp* from its ON state to OFF state, resulting in the formation of the final plasmid with both *attL/R* Int9 and *attL/R* Int13 sites. The corresponding colors and shapes indicate the genetic components of the switch.

### 2.2 *In vitro* function of Int-Plex@ binary memory switch system

Our first goal was to understand the behavior of this new binary memory system and ensure that the presence of *att* sites *in tandem* would not interfere with expected Int activity. For the functional evaluation, we chose cell-free *in vitro* transcription-translation reactions (TxTl), given its valuable application as a fast platform for testing Int activity, DNA editing tools and genetic circuits^35,36^. The switch was assembled using TxTl optimized parts, including the *degfp* gene as a reporter, Ribosome Binding Site (RBS) and the T7Max transcription system^35^ (Figure 2A). Reactions containing the reporter construction alone or with plasmids for Int expression were incubated overnight and analyzed for DNA rearrangement and deGFP fluorescence emission. This enabled us to test the efficacy of each reaction independently and transiently *in vitro* for the first time before integrating the memory switch into the plant genome. Besides deGFP expression, DNA excisions by BxB1 or phiC31 were detected, as predicted, by the PCR amplifications (Figure 2, B and C). The excised circular DNA molecule was confirmed by Sanger sequencing, where post-recombination *attR* site for BxB1 is present (Figure 3, D and E). Sanger sequencing of phiC31 treatments is in progress. All sequences are available for consultation at the link provided in the Data availability section. In addition, the introduction of Int9 or Int13 could invert the DNA and allowed deGFP production *in vitro* (Figure 3, A and B). Although the presence of both Int9 and Int13 in the same reaction must result in the re-establishment of the initial OFF state with *degfp* silencing, the closed nature of the reaction environment and relatively short incubation time preserved the protein produced while the single-edited intermediate was present, maintaining a high level of fluorescence. The double-editing event was confirmed by Sanger sequencing of the reporter construct, where both post-recombination *attL/attR* site pairs for Int9 and Int13 are present in the same molecule (Figure 3, C-E).

**Figure 2.**
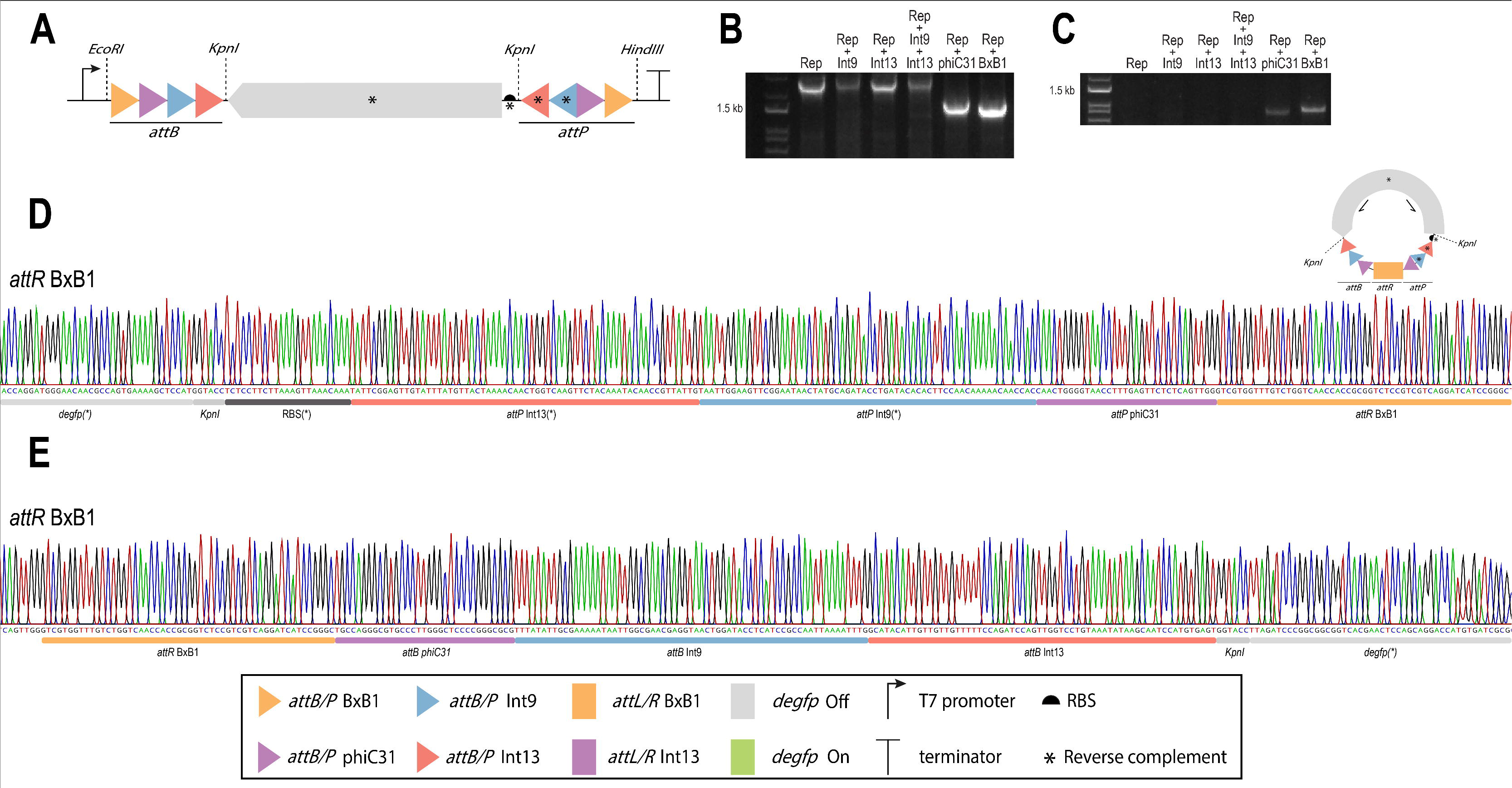
In vitro function of Int-Plex@ memory switch driven by serine integrases in cell-free TX-TL system. (A) Schematic overview of the IntPlex@ memory bimodular genetic switch architecture, which consists of a RBS sequence in reverse orientation (black) and a *degfp* sequence in reverse orientation (gray) surrounded by four *attB* and *attP* sites (triangles) of four serine integrases: BxB1 (yellow), phiC31 (lilac), Int9 (blue) and Int13 (red). (B) Agarose 1% gel of PCR reaction for each primer combination used to confirm the presence of edited DNA. (C) Agarose 1% gel of PCR reaction for each primer combination used to confirm the presence of the excised DNA with *attR* scar. (D and E) Sanger sequencing chromatograms of excised circular DNA with *attR* scar. The DNA fragments were obtained through PCR using pairs of primers indicated in Table 1. The position of the primers is indicated by black arrows. All sequences corresponded to the in silico predicted genome-edited sequences. The corresponding colors and shapes indicate the genetic components of the switch. First lane is 1 kb plus ladder (Life Technologies, USA). Reverse complementary sequences are indicated by a black asterisk. No off-target effects were detected in the analyzed sequences.

**Figure 3.**
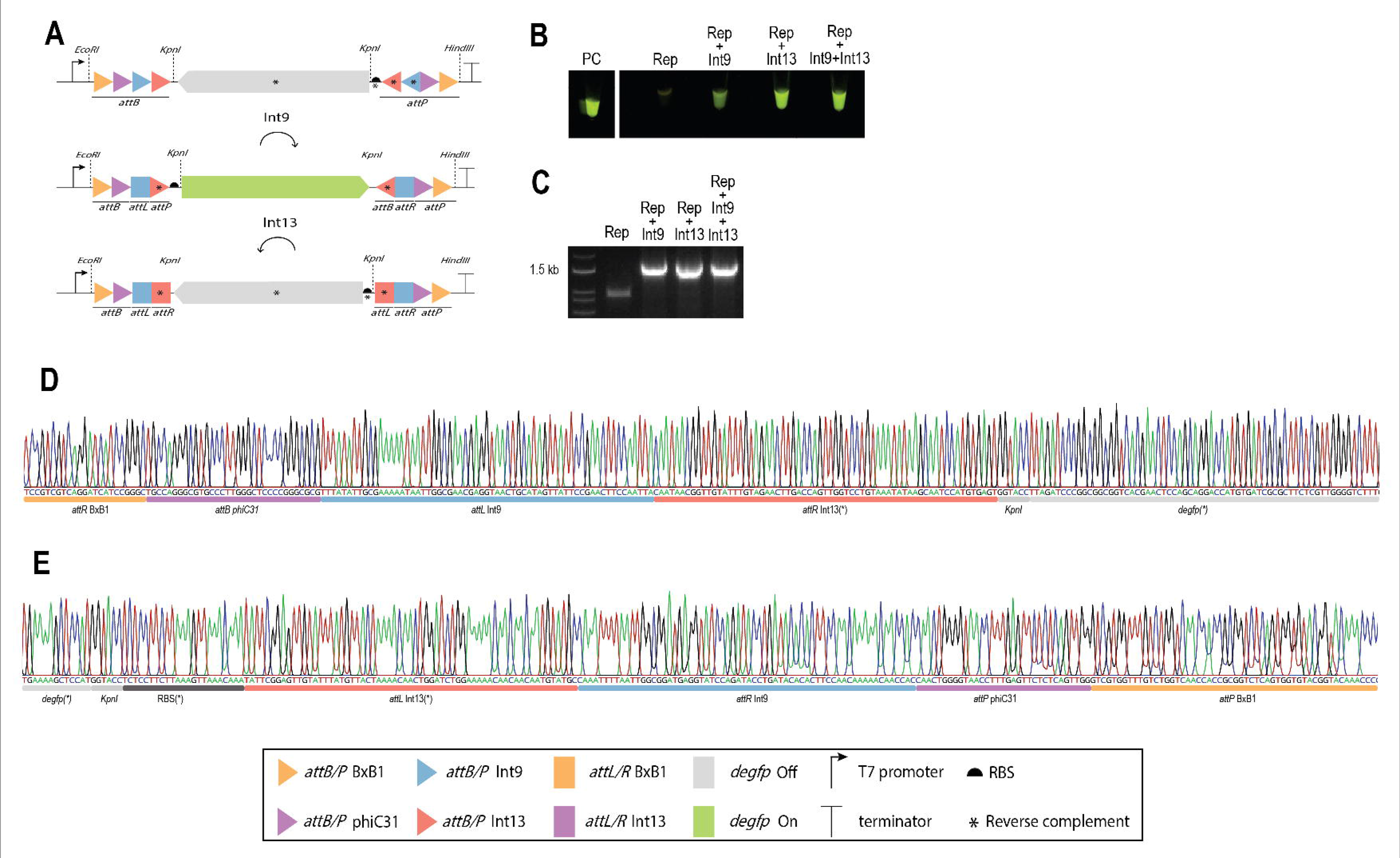
In vitro function of Int-Plex@ memory switch driven by serine integrases in cell-free TX-TL system. (A) The diagram shows the flipping of the *degfp* in two steps. In the first step, Int 9 recognizes its *attB/P* sites and catalyzes the inversion of *degfp* from its OFF state to ON state, resulting in the formation of the intermediate plasmid with *attL*/Int9 and *attR*/Int9 sites and the output deGFP. Instantaneously, this intermediate plasmid is the substrate for the action of Int13 that recognizes its *attB/P* sites and catalyzes the inversion of *degfp* from its ON state to OFF state, resulting in the formation of the final plasmid with both *attL/R* Int9 and sites *attL/R* Int13. (B) *In vitro* reactions after 24h post incubation of Int-Plex@ cassette plasmid with Int 9 and Int 13 plasmids in Arbor Biosciences™ myTXTL system. Furthermore, it is noteworthy that since both enzymes were added simultaneously (pART27+Int9+Int13), the ON state varies according to the first integrase to catalyze the plasmid inversion. Thus, the system presents the deGFP for several days according to the ON state mRNA molecules and protein half-lives. (C) Agarose 1% gel of PCR reaction for each primer combination used to confirm the presence of the inverted DNA with *attL/R* sites. (D and E) Sanger sequencing chromatograms of regions flanking the *degfp* obtained after simultaneous treatment with Integrases 9 and 13. The DNA fragments were obtained through PCR using pairs of primers indicated in Table 1. The position of the primers is indicated by black arrows. All sequences corresponded to the *in silico* predicted genome-edited sequences. The corresponding colors and shapes indicate the genetic components of the switch. First lane is 1 kb plus ladder (Life Technologies, USA). Reverse complementary sequences are indicated by a black asterisk. No off-target effects were detected in the analyzed sequences.

**Table 1.**
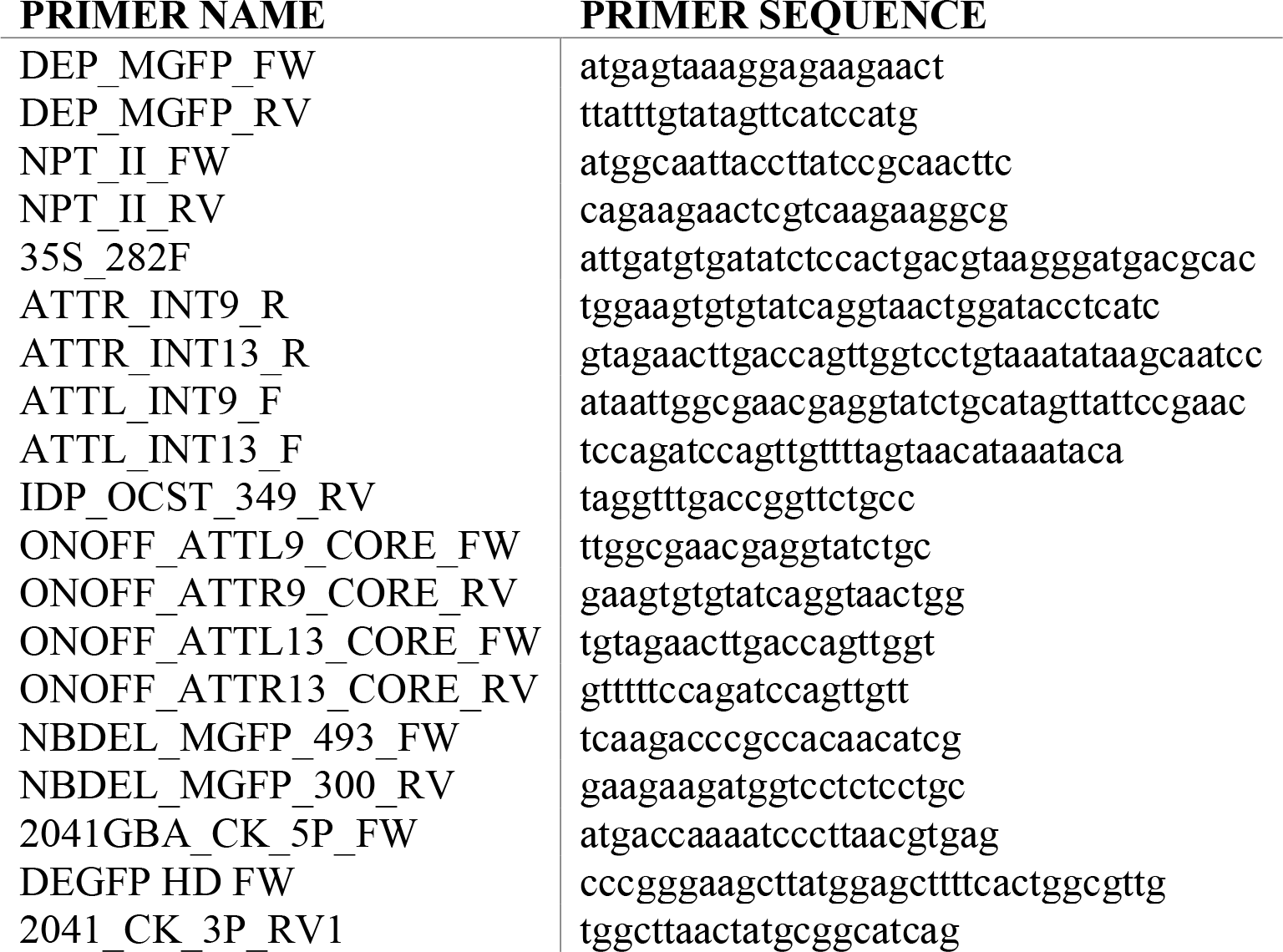

### 2.3 Int-Plex@ binary memory switch system design for plant genome modulation

To demonstrate the applicability of this tool in DNA modulation, we also verified the functionality of the switch stably integrated into the plant genome. Here, we assembled the reverse complement sequence of *mgfp* flanked by four in tandem BxB1, phiC31, Int9, and Int13 *attB* and *attP* sites in a synthetic genetic switch, denoted IntPlex@ binary memory switch and inserted into the *Nicotiana benthamiana* genome. Each serine integrase was plant codon optimized and inserted in a different set of plasmids for two systems of gene delivery: agroinfiltration by *Agrobacterium tumefaciens* and biolistic. The IntPlex@ binary memory switch takes these four serine integrase input signals to control an output (genome edition) in a user-defined manner (excision or inversion).

#### 2.3.1 Plant genome excision triggered by phiC31 or BxB1

BxB1 or phiC31 catalyzed the excision of *mgfp* from the genome. Upon delivery of the effector plasmids containing the integrases by agroinfiltration, molecular analyses of plant DNA within five d.p.i (days post infiltration) showed successful deletion, with genome sequencing confirmation of proper formation of respective *attL* sites for each Int (Figure 4A and Figure 5) and loss of DNA sequence flanked by its attachment sites (output) (Figure 4, B and C). Sanger sequencing of phiC31 treatments is in progress. Moreover, Nanopore sequencing was utilized to identify the excised circular DNA molecule containing *mgfp*, BxB1 *attR* site and remaining *attB/P* sites from the other integrases (Figure 6). The sequencing run lasted four hours, yielding roughly 800,000 reads. As depicted in Figure 7, the basecalled read lengths exhibited a bimodal distribution, with the median read length positioned within the distribution’s upper mode. Quality scores ranged along the x-axis, predominantly showing mid to high values indicative of reliable base calling accuracy. This pattern reflects a diverse set of read lengths, with shorter reads tending to have higher quality scores. For subsequent analysis, only reads with quality scores of 10 or above were maintained (383,245 reads). However, the detection of phiC31 output and stability and processing time for the circular DNA excised degradation remains to be evaluated. Regarding the delivery of the Int effector by biolistic, neither genome editing, nor the output molecule were detected (data not shown). Nanopore sequencing will be applied to detect the output molecules and analyze the low detection rate. All sequences are available for consultation at the link provided in the Data availability section.

**Figure 4.**
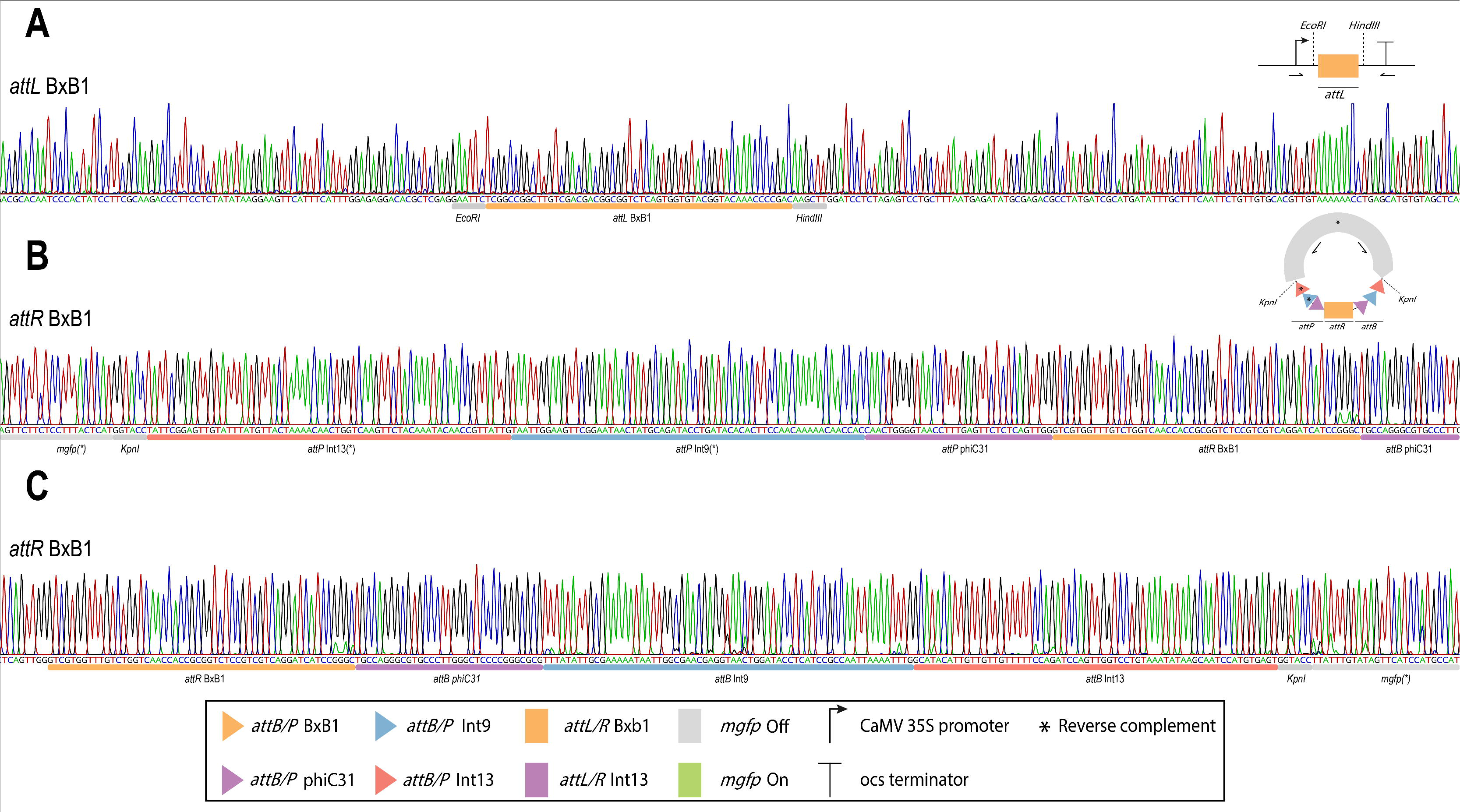
IntPlex@ memory genetic switch for DNA excision driven by BxB1. (A) Sanger sequencing chromatogram of edited plant genome with attL/BxB1 scar. (B and C) Sanger sequencing chromatograms of excised circular DNA with *attR* scar. The DNA fragments were obtained through PCR using pairs of primers indicated in Table 1. The position of the primers is indicated by black arrows. All sequences corresponded to the in silico predicted genome-edited sequences. The corresponding colors and shapes indicate the genetic components of the switch.

**Figure 5.**
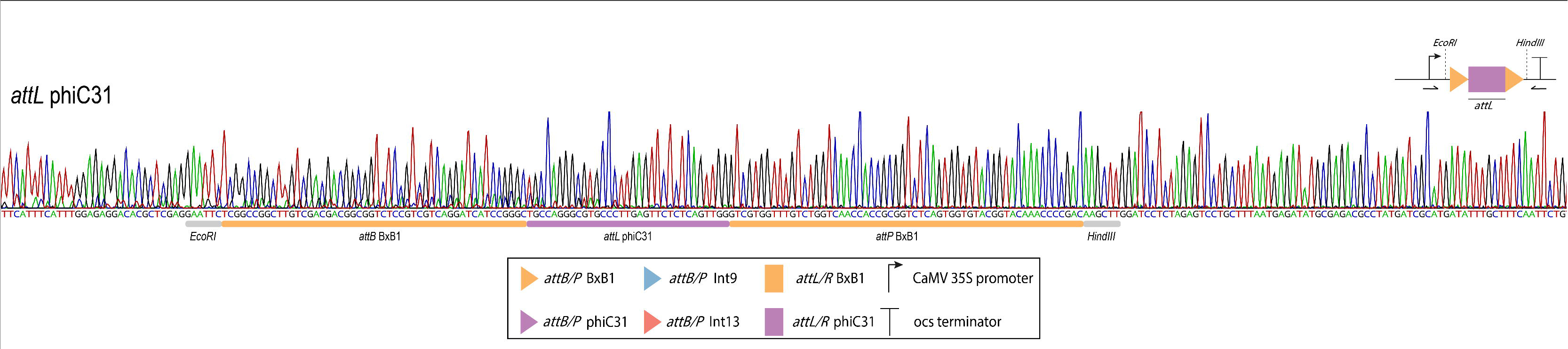
IntPlex@ memory genetic switch for DNA excision driven by phiC31. Sanger sequencing chromatogram of edited plant genome with phiC31 *attL*. The DNA fragments were obtained through PCR using pairs of primers indicated in Table 1. The position of the primers is indicated by black arrows. The sequence corresponded to the in silico predicted genome-edited sequence. The corresponding colors and shapes indicate the genetic components of the switch.

**Figure 6.**
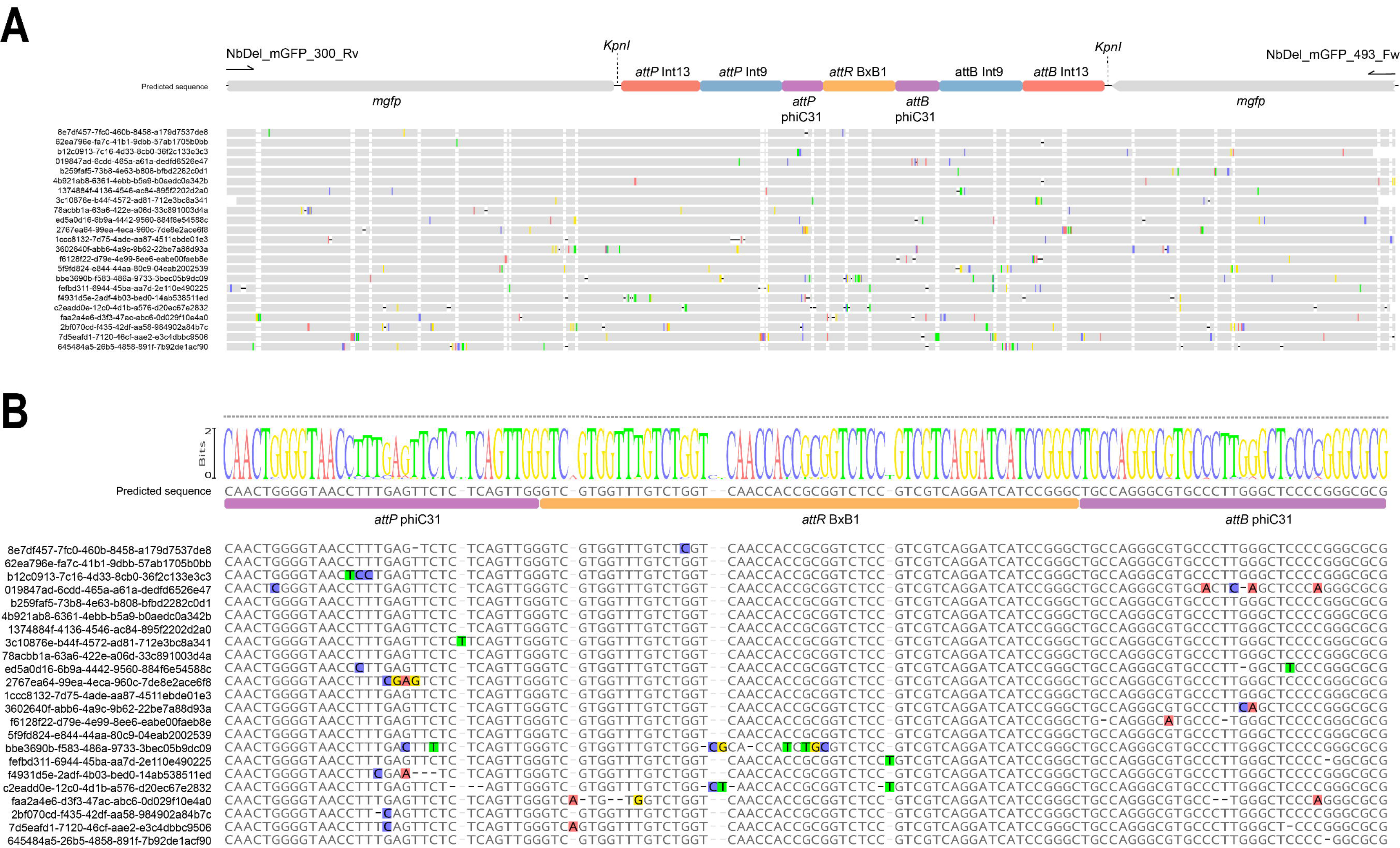
Nanopore sequencing evidence of genetic switch operation driven by BxB1. (A) The diagram illustrates the nanopore raw reads mapped to the predicted excised circular DNA molecule with BxB1 *attR*. The corresponding colors and shapes indicate the genetic components of the switch. (B) A detailed view of the excision region, showing the resulting BxB1 *attR* and the flanking phiC31 *attP/attB* sites.

**Figure 7.**
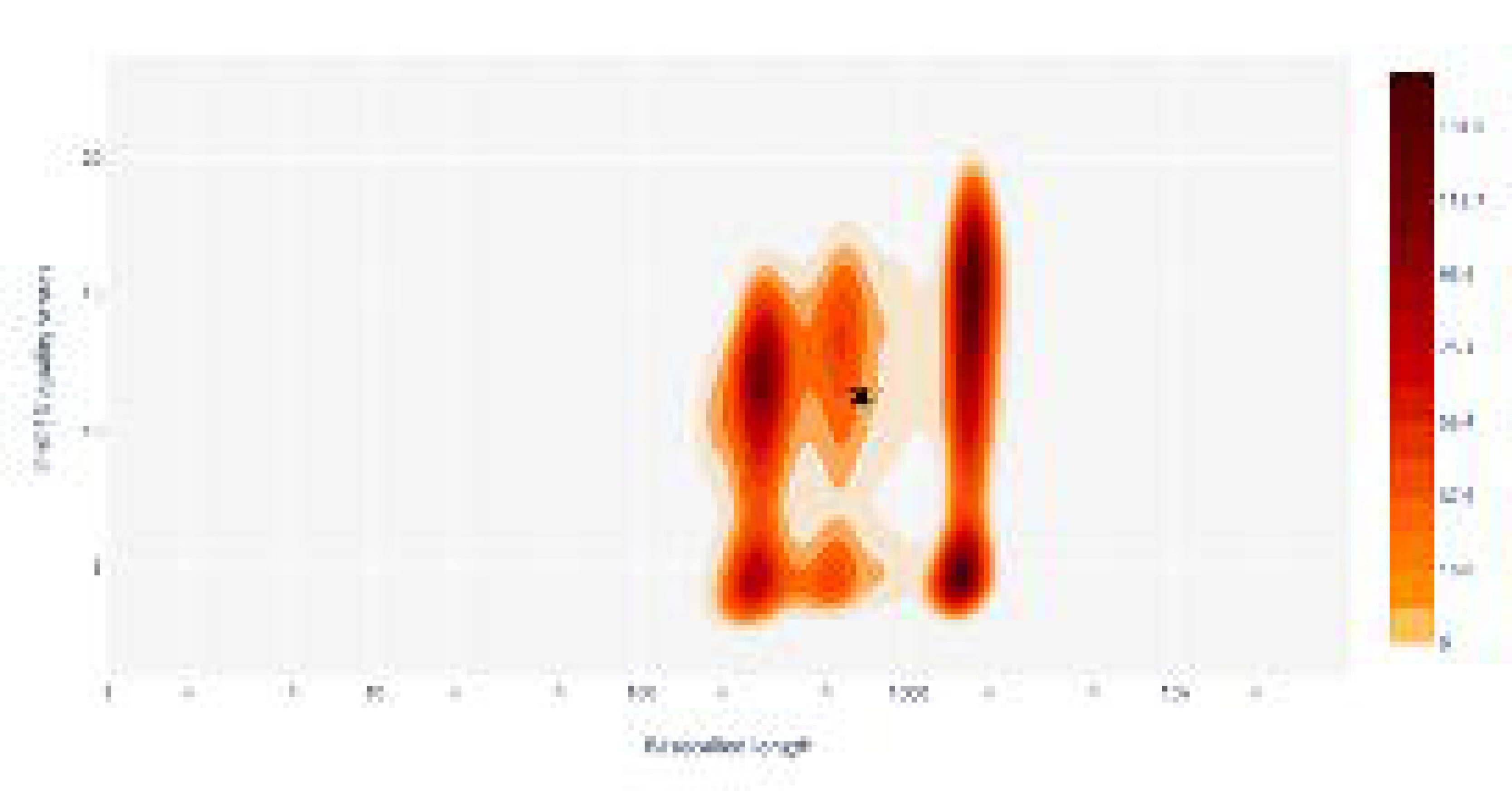
Basecalled reads length vs reads quality score. The color gradient indicates the density of reads, with warmer colors representing higher densities.

#### 2.3.2 Plant genome inversion triggered by Int 9 or Int 13

This switch operates as an ON switch, where MGFP is expressed in the ON state and not in the OFF state. Upon delivery of the effector plasmids containing the integrases by agroinfiltration, molecular analyses of plant DNA within five d.p.i showed successful DNA inversion. Int9 recognized its *attB/P* sites and catalyzed the inversion of *mgfp* from its OFF state to ON state, resulting in the formation of the intermediate genome with *attL*/Int9 and *attR*/Int9 sites and the output MGFP after int delivery via agroinfiltration (Figure 8A). Likewise, Int13 recognized its *attB/P* sites and catalyzed the inversion of *mgfp* from its OFF state to ON state, resulting in the formation of the intermediate genome with *attL*/Int13 and *attR*/Int13 sites and the output MGFP (Figure 8B). As for effector Int delivery by biolistic, phenotypical analysis showed no increase in mGFP fluorescence (data not shown). However, fluorescence could be detected in the positive control of plants bombarded with microparticles coated with a plasmid constitutively expressing eGFP^37^. Despite the lack of signal, DNA inversion could be detected for both Ints by PCR amplification, with confirmation of inversion and recombined *attL/R* sites by DNA sequencing.

**Figure 8.**
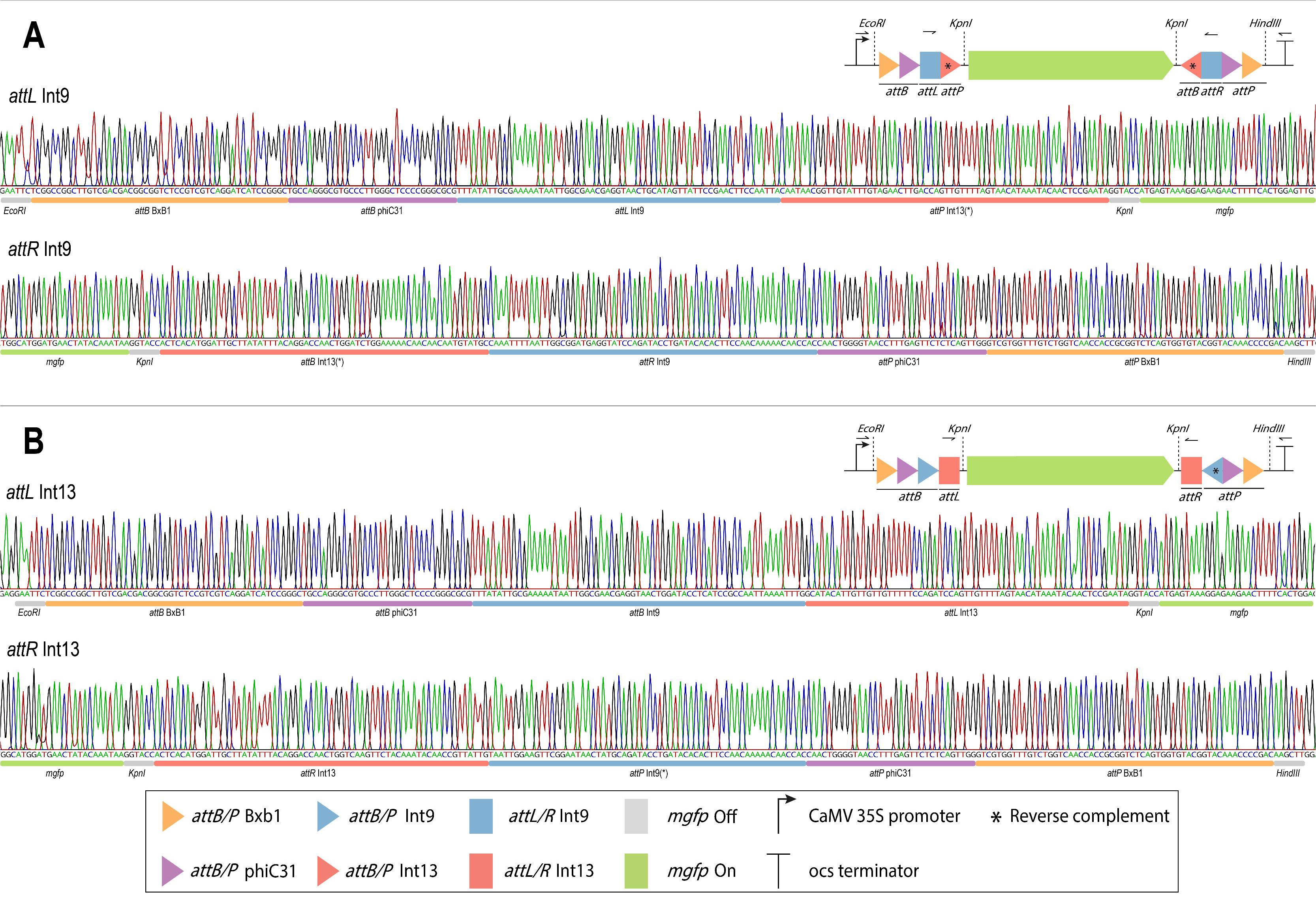
IntPlex@ memory genetic switch for gene modulation (turn gene on) driven by Int9 or Int13. (A) Sanger sequencing chromatogram of edited plant genome with Int9 *attL* and *attR* post recombination sites. The sequence shows the flipping of the DNA flanked by the Int9 *attB* and attP sites and, consequently, the inversion of the *mgfp* to its coding sequence (MGFP output). The Int13 *att* sites are also inverted and oriented towards their reverse complement. (B) Sanger sequencing chromatogram of edited plant genome with Int13 *attL* and *attR* post recombination sites and the inversion of the *mgfp* to its coding sequence (MGFP output). The fragments were obtained through PCR using pairs of primers indicated in table 1. The position of the primers is indicated by black arrows. A pair of primers was used to amplify the complete *attL* site (black upper arrow) and a second pair of primers to amplify the *attR* site (black lower arrow). The sequences corresponded to the in silico predicted genome-edited sequences. The corresponding colors and shapes indicate the genetic components of the switch.

#### 2.3.3 Plant genome modulation: Turning DNA ON and OFF

While a one-way memory switch is functional, reversibly turning a gene on and off would allow interesting bidirectional regulation. Thus, Int9 and Int13 were agroinfiltrated into the plant five days apart to turn *mgfp* on and off in the same cell. Initially, Int 9 recognized its *attB/P* sites and catalyzed the inversion of *mgfp* from its OFF state to ON state, resulting in the formation of the intermediate genome with *attL*/Int9 and *attR*/Int9 sites and the output MGFP. Five days after, this intermediate genome was the substrate for the action of Int13 that recognized its *attB*/P sites and catalyzed the inversion of *mgfp* from its ON state to OFF state, resulting in the formation of the final genome with both *attL*/Int9 and *attR*/Int9 and *attL*/Int13 and *attR*/Int13 sites. Thus, the system presents the MGFP until the ON state mRNA molecule can no longer be translated according to the MGFP half-life time. The Sanger sequencing of Int9 and Int13 treatments are in progress. All sequences are available for consultation at the link provided in the Data availability section.

## 3. DISCUSSION

In this study, our primary goal was to broaden the plant genome modulation toolbox by creating a binary memory switch controlled by serine integrases. Currently, the available array of tools for constructing synthetic memory systems in plants is quite limited, mainly relying on phiC31 and BxB1^29,31,32,38,39^. Efforts for the systematic identification of new Ints and their *att* sites, including metagenomic studies, development of new bioinformatic approaches, and functional characterization, contribute to expanding the availability of such tools and enhance the possibilities for assembly of more complex multicomponent genetic circuits^40^. In that context, we applied phiC31, BxB1, and two more recent Ints (Int9 and Int13) previously characterized in a plant model to create a bimodular switch for DNA excision and bidirectional inversion^25^. All four Int successfully rearranged the target DNA inserted into the genome of *N. benthamiana* as programmed, confirming the functionality of both modules in our system.

More than assembling a new switch system for gene expression regulation, this proof-of-concept demonstration paves the way for applying new Int candidates other than phiC31 and BxB1 in plants. Beyond orthogonality, which allows for various effectors in the same circuit, different integrases can show a range of efficiency degrees beneficial for implementing flow control nodes in a synthetic regulatory pathway. As demonstrated by Gomide et al. (2020), for instance, Int13 and Int5 both could invert a target DNA sequence in *Arabidopsis thaliana* protoplasts, although resulting in different levels of reporter expression^25^.

Another essential advantage point of IntPlex@ is the use of independent Ints to control inversion directionality. After the first inversion event by Int9 or Int13, restoration to the initial OFF state depends exclusively on introducing the second enzyme, with no chance of interference. In contrast, the RESET system published by Bernabé-Orts et al. (2020) applies phiC31 with its correspondent RDF protein to revert unidirectionality^29^. Although aiming at somewhat different outcomes, it is noteworthy that with RDF, they observed an increase in system instability. Recombination of *attL/R* sites requires direct interaction of Int-RDF, but the presence of free phiC31 led to new recombination of the *attB/P* sites just restored, resulting in mixed states of the switch at the same time^29^.

The Sanger Sequencing method is widely used for identifying plant genome editions. Nevertheless, it comes with some limitations, including the prerequisite for pre-existing knowledge of the DNA sequence and a constrained ability to simultaneously detect various DNA fragments. On the other hand, the utilization of Nanopore DNA sequencing has enabled determination of total DNA products of the plant cells genome edition reactions in a short time and with high-throughput data analysis in real time. Nanopore sequencing holds great promise for detecting edited DNA and when combined with stoichiometry analysis, this technology provides a powerful toolset to access genomic diversity for studying edited DNA at a quantitative level. The integration of nanopore sequencing and stoichiometry analysis not only enhances the ability to detect edits accurately but also facilitates a deeper understanding of the dynamics and efficiency of serine integrase editing reactions.

It is worth mentioning that when a system is imported from the prokaryotic to the eukaryotic cell, as is the case for bacteriophage-derived recombinases, it is necessary to consider the presence of the nucleus as a physical barrier to the performance of enzymes that act on DNA. Nevertheless, all four integrases tested in this work were able to overcome the nuclear barrier and interact with chromatin without the need to add NLS or other protein engineering. It is possible that the passage through the nuclear pore occurred due to the presence of intrinsic NLS in its native structure after exposure of positively charged amino acids during protein folding^25^. Notably, the system functioned consistently over two plant generations. Employing large serine-integrases in modular switch devices exemplifies a sophisticated and controlled approach, affording precision in genetic alterations with the potential for intricate modulation of plant physiological processes. Consequently, the integration of large serine-integrases into the biotechnological toolkit heralds a promising trajectory for refining and expanding strategies in plant genetic engineering. However, further research is needed to fully understand the potential of this system in plant biotechnology, the potential for unintended off-target effects, and to optimize its use.

## 4. CONCLUSIONS

PhiC31, BxB1, Int9, and Int13 promoted efficient directional DNA site-specific recombination of complex chromatin of the plant cell, and thus have major potential applications in genome engineering and metabolic pathway engineering. This memory genetic switch system is dynamic and may be combined with other technologies like inducible promoters and advanced sequencing tools. For example, implementing the sequence barcoding with Nanopore sequencing amplifies the capabilities of genomic analysis and allows for comprehensive profiling of the entire edited genome, capturing structural variations or ensuring the fidelity of the editing process. Together, this combined approach not only validates the accuracy of serine integrase-mediated edits at a detailed level but also offers a holistic view of the entire genome modulation, fostering a deeper understanding of the genomic landscape post-editing. Lastly, it is more than evident that we are living in an era marked by the development of tools that allow crop genomes to be edited to provide resistance to environmental challenges, improve nutritional content and optimize defenses against pathogens. However, the lack of efficacy of plasmid delivery systems has hampered advancements in the precision of gene editing. This is a crucial moment that requires the convergence of efforts to develop studies that aim to refine plasmid or gene delivery systems, ensuring that genetic modifications reach the maximum number of cells efficiently.

## 5. MATERIAL AND METHODS

### 5.1 mGFP genetic switch plasmid design

In pLSB_pCAMBIA2300_IntX or pUC_IntX plasmids, both *actin2* gene promoter and *NOS* terminator drive (a) plant codon-optimized serine integrases transcription for heterologous expression or (b) serine integrases fused with NLS signal. In pLSB_PVX_GW_IntX plasmids, both subgenomic CP promoter and *NOS* terminator drive serine integrase transcription for heterologous expression with or without the NLS signal.

### 5.2 Serine-Integrase Plasmid Design

The *Int*9, *Int13*, *phiC31*, and *BxB1* genes were retrieved from pUC57^25^ and cloned into PVX-GW binary plasmid^37^ by Epoch Life Science Inc. These plasmids resulted in a set of serine integrase expression plasmids, individually called pLSB_PVX_GW_X (X = *Int9*, *Int13*, *phiC31*, or *BxB1*). The *Int9, Int13, phiC31,* and *BxB1* genes controlled by the *actin2* gene promoter and *NOS* terminator were retrieved from pUC57 and cloned in pCAMBIA2300 plasmid by Epoch Life Science Inc^25^. These plasmids resulted in a set of serine integrase expression plasmids, individually called pLSB_pCAMBIA2300_X (X = *Int9*, *Int13*, *phiC31*, or *BxB1*).

### 5.3 Plant transformation

Transgenic plants of *Nicotiana benthamiana* were produced at University of California Riversidés Department of Botany and Plant Science performed the plant transformation. pART27 plasmid containing the genes BXB1_PHIC31_9_13_mGFP(rc) and nptII were delivered into GV3101 *Agrobacterium tumefaciens* via electroporation. Antibiotic-resistant colonies were selected and confirmed by colony PCR for glycerol stocks. The transformed *A. tumefaciens* strain was used to inoculate the leaf tissue of six-week-old plants. Callus induction and shoot induction media containing MS salts, MS vitamins, 0.56 0.56mM Myo, 8.84 8.84 μM BAP, 0.54 0.54 μM NAA, 3% sucrose, 0.21 0.21 mM kanamycin, 0.55 0.55 mM Cefatoxime and 1.321.32 mM Carbenecillin were used to induce calli and shoots. Thirty and ten shoots transformed with BXB1_PHIC31_9_13_mGFP(rc) and pART27-nptII (empty plasmid) respectively were transferred to rooting media containing ½ MS salts, MS vitamins, 0.56 0.56 mM Myo-inositol, 0.98 0.98 μM IBA, 1.5% sucrose, 0.34 0.34 mM vancomycin, 0.55 0.55 mM cefotaxime, and 1.32 mM Carbenicillin. A total of 40 *N. benthamiana* plants were analyzed by end-PCR using specific primers for each construct. Twenty-one plants amplified an expected band of 717 bp with the specific primers FwmGFP/RvmGFP, and nine plants transformed with pART27 (empty plasmid) amplified a band of 699 bp for the nptII gene. *Nicotiana benthamiana* transgenic seeds germinated in vermiculite, and plantlets were grown in a greenhouse. Eight-week-old plants were used for the Agrobacterium infiltration or biolistic procedure.

### 5.4 *In vitro* Transcription/Translation Reactions

*In vitro*, Transcription/Translation Reactions, or TxTl reactions, were performed with myTxTl T7 Expression Kit (Daicel Arbor Biosciences) following manufacturer instructions. Briefly, 24 μL reactions were assembled in 1.5 mL centrifugation tubes with 18 μL Sigma 70 Master Mix, 0.1 nM P70a-T7rnap HP, and 10 nM of each plasmid (reporter + effector Int). Reactions were incubated at 29°C for 16 h. After incubation, GFP production was visually assessed with an LED blue light Transilluminator (KASVI), and 2 μL were used as a template for PCR amplification.

### 5.5 Delivery Systems

#### 5.5.1 Agroinfiltration

The binary plasmids were electroporated into *Agrobacterium tumefaciens* strain GV3101 and selected on LB agar supplemented supplemented with 0.1 mM kanamycin and 0.1 mM gentamicin.. After two days of growth, several colonies were carefully harvested from the plate and suspended in LB medium (2 mL). The bacteria were grown overnight on an orbital shaker at 28 °C at 200 rpm. The cells were collected by centrifugation for 15 min at 4000 g, and the precipitate was resuspended in the 1x MMA infiltration buffer (10 mM MES, pH = 5.6, 10 mM MgCl_2_, 180 µM acetosyringone) to OD600 = 0.3. Agrobacteria harboring plasmids with each integrase and 35S: p19 gene were mixed in equal volumes. Leaves of 8-week-old *N. benthamiana* plants were injected with agrobacteria suspension using a syringe without a needle. Agroinfiltration assays were assessed with and without co-infiltration with 35S: p19 with no changes in results. The infiltrations were performed on the same plant, and duplicate infiltrations were performed on at least three plants within a single experimental assay.

#### 5.5.2 Biolistic

Genetic modified *N. benthamiana* plant leaves carrying inverted *mgfp* gene flanked by integrases attachment sites were directly shot with gold (1.5-3.0 µm) and tungsten (approx. 1µm) microparticles as previously described^41^. The microparticles were coated with pUC27 plasmids carrying *Int9*, *Int13*, *phiC31*, and *BxB1* integrase genes individually^25^. As a positive control of gene delivery, a plasmid carrying a constitutive promoter controlling GFP gene was shot under the same conditions at wild-*type N. benthamiana* leaves. Microparticles mixed with 8 μL DNA (1μg.μL^-1^) were accelerated at 650 psi into plant leaves (2,5-3,5 cm length) at a 2 cm distance. Eight shots of each DNA construction were bombarded in 8-week-old plants. 72h after foreign DNA was delivered, shooted leaves were analyzed on Confocal laser scanning microscope (Leica TCS SP8, GER) to visualize *mgfp* expression. PCR screening has been performed to detect *mgfp* and the *attL* and *attR* post-recombination sites.

### 5.6 Plant total DNA extraction and PCR

Total plant DNA isolation was based on the modified CTAB Method Protocol described by Lacorte et al. (2010) with adaptations^39^. Firstly, plants were screened by PCR using primer pairs DEP_mGFP_FW and DEP_mGFP_RV - designed to detect *mgfp* – and nptII_Fw and npt_II_Rv - for the amplification of *nptII*, a kanamycin resistance marker – to identify positive transformants. Following the integrase delivery method and incubation period, we collected slices of approximately 20mg from leaves of treated and control plants and proceeded with DNA extraction, as mentioned above. DNA samples were then used as templates in PCR reactions for detection of either *attL* and *attR* sites resulting from *Int9* or *Int13* recombination or the deletion of *mgfp* gene flanked by *phiC31* and *BxB1 att* sites upon introduction of these integrases in the system. For *attL* and *attR* sites amplification, we have used the primer combinations 35S_282F + attR_IntX_R and attL_IntX_F + IDP_OCSt_349_Rv, respectively, with “IntX” on the primer name denoting the use of specific primers for each Int. As for deletion confirmation, primer pair 35S_282F + IDP_OCSt_349_Rv was used to amplify the whole cassette between the 35S promoter and *OCS* terminator, with a band shift of about 1kb indicating *mgfp* excision. The circular molecule containing *the mgfp* gene excised from the system by phiC31 or BxB1 was amplified using the primer pair NbDel_mGFP_493_Fw + NbDel_mGFP_300_Rv. Two PCR reactions were necessary for the biolistic method to amplify *attL* and *attR* sites. The primer pair 35S_282F + IDP_OCST_349_RV performed the first round, which amplified the cassette. The first reaction was then used as a template for the specific reaction with primers attached on the *attL* and *attR* sites, paired with terminator and promoter primers, respectively. That way, the second round of amplification for detection of the Int9 *attL* site used primers 35S_282F + OnOff_attR9_core_Rv, for Int9 *attR* site primers OnOff_attL9_core_Fw + IDP_OCST_349_RV, for Int13 *attL* site primers 35S_282F + OnOff_attR13_core_Rv and to amplify Int13 *attR* site primers OnOff_attL13_core_Fw + IDP_OCST_349_RV. As for the primers used for molecular detection in TxTl reactions, inversion of the reporter was amplified with primer pair 2041GbA_ck_5p_FW + deGFP_Hd_Fw while excision was detected with primers 2041GbA_ck_5p_FW and 2041_ck_3p_RV1. Primers used in this study are listed in Table 1.

Amplification reactions were performed in a Veriti™ 96-Well Fast Thermal Cycler (Applied Biosystems, USA). Reaction set-up included 2.5ul of 10X PCR Buffer (Thermo Scientific, USA), 1.5 mM MgCl2, 0.4 mM dNTP mix, 1 U Platinum™ Taq DNA Polymerase (5 U/ μL) (Thermo Scientific, USA), 0.2 μM each primer and 1 μL of purified DNA in a 25 μL final volume. PCR cycling consisted of one initial incubation at 94°C for 2 minutes, followed by 40 cycles of denaturation at 94°C, 30s; Annealing TEMP for 30s; Extension at 72°C for 1 min/kb, with final incubation at 4°C. Given that primer pair 35S_282F + IDP_OCSt_349_Rv will amplify both intact and disrupted cassette resulting from *mgfp* gene deletion by phiC31 or BxB1, an extension time of just 30 seconds were used in this case to favor amplification of the 1kb smaller amplicon. All PCR products were detected in 1% agarose gel electrophoresis stained with SYBR™ Safe DNA Gel Stain (Invitrogen, USA). Gel slices containing the specific bands were excised under LED Blue Light transluminator (KASVI, Taiwan) to prevent nicking of the DNA and its purification was carried out using the ReliaPrep™ DNA Clean-Up and Concentration System (Promega, USA) according to manufacturer specifications. Isolated amplicons were cloned in pGEM-T-Easy (Promega, USA) and sequenced (Macrogen, USA) after DNA propagation in *Escherichia coli* DH10β strain.

### 5.7 Confocal laser scanning Microscopy

Leaf samples were analyzed using the Confocal laser scanning microscope (Leica TCS SP8, GER) after 5 d.p.i. Argon laser line excitation wavelength and emission bandpass filter wave lengths for MGFP were 484 nm and 500-550 nm. Chlorophyll autofluorescence was detected, in parallel with MGFP acquisition, using a 650–710 nm bandpass filter. Image acquisition parameters (e.g. laser power, pinhole, detector gain, etc.) and sampling time post-infiltration were held constant within an experiment (i.e. within each figure). Raw data were processed in the LAS X (Leica Application Suite, GER) software (www.leica-microsystems.com, GER).

### 5.8 Nanopore sequencing and Data analysis

Library preparation was carried out using the ligation sequencing kit (Oxford Nanopore Technologies, UK) SQK-NBD112-24, according to the manufacturer’s instructions. Briefly, all amplicons were purified, and approximately 200 fmol per sample were submitted to end-prep and native barcode ligation, using NEBNext Ultra II End repair/dA-tailing Module (New England Biolabs, USA) and NEB Blunt/TA Ligase Master Mix (New England Biolabs, USA) respectively. Next, native adapter ligation and cleaning steps were performed using NEBNext Quick Ligation Module (New England Biolabs, USA) kit and Agencourt AMPure XP beads (Beckman Coulter, USA), respectively. The library was sequenced on a R10.4 flow cell (FLO-MIN112) (Oxford Nanopore Technologies, UK) on a MinION Mk1b sequencer with MinKNOW (23.04.6). Basecalling, demultiplexing and adapter/barcode trimming were performed using Guppy [v6.5.7]^42^, with a Super accurate (SUP) basecalling model. The reads with barcodes on both ends and quality scores ≥ 10 were mapped against all the in silico predicted genome-edited sequences, including the predicted circular molecules containing the excised *mgfp* gene, using minimap2^43^.

## Acknowledgements

We acknowledge funding support from Embrapa Genetic Resources and Biotechnology/National Institute of Science and Technology in Synthetic Biology, National Council for Scientific and Technological Development/ Ministry of Agriculture Livestock and Supply (465603/2014-9; 400145/2023-5), Research Support Foundation of the Federal District (0193.001.262/2017), and Coordination for the Improvement of Higher Education Personnel. We thank Plant Transformation Research Center, University of California-Riverside, USA, for producing the transgenic plants of *N. benthamiana*; Plant Germplasm Quarantine Station, Embrapa Genetic Resources and Biotechnology, Brazil, for regulating the importation of *N. benthamiana* (21016.000015/2020-21; 21016.004393/2020-83).

## Author contributions

M.A.O.: conceptualization, investigation, data curation, and writing - original draft. R.N.L.: conceptualization, investigation, data curation, and writing - original draft. L.H.F.: investigation, data curation, and analysis. M.M.S.A.: investigation, data curation, and analysis. F.L.M.: investigation, data curation, and analysis. R.V.B.: investigation, data curation, and analysis. C.L.: investigation, data curation, and analysis. E.L.R.: funding acquisition, supervision, project administration, and writing - review & editing. All authors revised the manuscript and approved the final version.

## Data availability

The sequencing data obtained in this study are available in: https://github.com/Rech-PBSyn/Serine_Integrases

## Additional information

Additional Supporting Information may be found in the online version of this article at the publisher’s website or under request. Correspondence and requests for materials should be addressed to Elibio Rech.

## REFERENCES

1 Ortega, C., Abreu, C., Oppezzo, P. & Correa, A. Overview of High-Throughput Cloning Methods for the Post-genomic Era. *Methods in molecular biology (Clifton*, N.J.) 2025, 3–32, doi:10.1007/978-1-4939-9624-7_1 (2019).

2 Casini, A., Storch, M., Baldwin, G. S. & Ellis, T. Bricks and blueprints: methods and standards for DNA assembly. Nature reviews. Molecular cell biology 16, 568–576, doi:10.1038/nrm4014 (2015).

3 Gibson, D. G. et al. Enzymatic assembly of DNA molecules up to several hundred kilobases. Nature methods 6, 343–345, doi:10.1038/nmeth.1318 (2009).

4 Gibson, D. G. Enzymatic assembly of overlapping DNA fragments. Methods in enzymology 498, 349–361, doi:10.1016/b978-0-12-385120-8.00015-2 (2011).

5 Sleight, S. C., Bartley, B. A., Lieviant, J. A. & Sauro, H. M. In-Fusion BioBrick assembly and re-engineering. Nucleic acids research 38, 2624–2636, doi:10.1093/nar/gkq179 (2010).

6 Vazquez-Vilar, M. et al. GB3.0: a platform for plant bio-design that connects functional DNA elements with associated biological data. Nucleic acids research 45, 2196–2209, doi:10.1093/nar/gkw1326 (2017).

7 Arber, W. Polylysogeny for bacteriophage lambda. Virology 11, 250–272, 10.1016/0042-6822(60)90065-9 (1960).

8 Mahmood, M. A., Naqvi, R. Z., Rahman, S. U., Amin, I. & Mansoor, S. Plant Virus-Derived Vectors for Plant Genome Engineering. Viruses 15, doi:10.3390/v15020531 (2023).

9 Khakhar, A. & Voytas, D. F. RNA Viral Vectors for Accelerating Plant Synthetic Biology. Frontiers in plant science 12, 668580, doi:10.3389/fpls.2021.668580 (2021).

10 Zulfiqar, S. et al. Virus-Induced Gene Silencing (VIGS): A Powerful Tool for Crop Improvement and Its Advancement towards Epigenetics. International journal of molecular sciences 24, doi:10.3390/ijms24065608 (2023).

11 Jagram, N. & Dasgupta, I. Principles and practice of virus induced gene silencing for functional genomics in plants. Virus genes 59, 173–187, doi:10.1007/s11262-022-01941-5 (2023).

12 Rajput, M. et al. RNA Interference and CRISPR/Cas Gene Editing for Crop Improvement: Paradigm Shift towards Sustainable Agriculture. *Plants (Basel*, Switzerland) 10, doi:10.3390/plants10091914 (2021).

13 Zhao, J. H. & Guo, H. S. RNA silencing: From discovery and elucidation to application and perspectives. Journal of integrative plant biology 64, 476–498, doi:10.1111/jipb.13213 (2022).

14 Jin, L., Chen, M., Xiang, M. & Guo, Z. RNAi-Based Antiviral Innate Immunity in Plants. Viruses 14, doi:10.3390/v14020432 (2022).

15 Metje-Sprink, J., Menz, J., Modrzejewski, D. & Sprink, T. DNA-Free Genome Editing: Past, Present and Future. Frontiers in plant science 9, 1957, doi:10.3389/fpls.2018.01957 (2018).

16 Pandey, P. K. et al. Versatile and multifaceted CRISPR/Cas gene editing tool for plant research. Seminars in cell & developmental biology 96, 107–114, doi:10.1016/j.semcdb.2019.04.012 (2019).

17 Adli, M. The CRISPR tool kit for genome editing and beyond. Nature communications 9, 1911, doi:10.1038/s41467-018-04252-2 (2018).

18 Lin, Q. et al. Genome editing in plants with MAD7 nuclease. Journal of genetics and genomics = Yi chuan xue bao 48, 444–451, doi:10.1016/j.jgg.2021.04.003 (2021).

19 Smith, M. C. M. Phage-encoded Serine Integrases and Other Large Serine Recombinases. Microbiology spectrum 3, doi:10.1128/microbiolspec.MDNA3-0059-2014 (2015).

20 Yang, L. et al. Permanent genetic memory with >1-byte capacity. Nature methods 11, 1261–1266, doi:10.1038/nmeth.3147 (2014).

21 Olorunniji, F. J. et al. Multipart DNA Assembly Using Site-Specific Recombinases from the Large Serine Integrase Family. *Methods in molecular biology (Clifton*, N.J.) 1642, 303–323, doi:10.1007/978-1-4939-7169-5_19 (2017).

22 Wang, X. et al. Bxb1 integrase serves as a highly efficient DNA recombinase in rapid metabolite pathway assembly. Acta biochimica et biophysica Sinica 49, 44–50, doi:10.1093/abbs/gmw115 (2017).

23 Andres, J., Blomeier, T. & Zurbriggen, M. D. Synthetic Switches and Regulatory Circuits in Plants. Plant physiology 179, 862–884, doi:10.1104/pp.18.01362 (2019).

24 Kassaw, T. K., Donayre-Torres, A. J., Antunes, M. S., Morey, K. J. & Medford, J. I. Engineering synthetic regulatory circuits in plants. Plant science : an international journal of experimental plant biology 273, 13–22, doi:10.1016/j.plantsci.2018.04.005 (2018).

25 Gomide, M. S. et al. Genetic switches designed for eukaryotic cells and controlled by serine integrases. Communications Biology 3, 255, doi:10.1038/s42003-020-0971-8 (2020).

26 Stark, W. M. Making serine integrases work for us. Current opinion in microbiology 38, 130–136, doi:10.1016/j.mib.2017.04.006 (2017).

27 Lloyd, J. P. B. et al. Synthetic memory circuits for stable cell reprogramming in plants. Nature biotechnology 40, 1862–1872, doi:10.1038/s41587-022-01383-2 (2022).

28 Bonnet, J., Subsoontorn, P. & Endy, D. Rewritable digital data storage in live cells via engineered control of recombination directionality. Proceedings of the National Academy of Sciences of the United States of America 109, 8884–8889, doi:10.1073/pnas.1202344109 (2012).

29 Bernabé-Orts, J. M. et al. A memory switch for plant synthetic biology based on the phage ϕC31 integration system. Nucleic acids research 48, 3379–3394, doi:10.1093/nar/gkaa104 (2020).

30 Merrick, C. A., Zhao, J. & Rosser, S. J. Serine Integrases: Advancing Synthetic Biology. ACS synthetic biology 7, 299–310, doi:10.1021/acssynbio.7b00308 (2018).

31 Thomson, J. G. et al. The Bxb1 recombination system demonstrates heritable transmission of site-specific excision in Arabidopsis. BMC biotechnology 12, 9, doi:10.1186/1472-6750-12-9 (2012).

32 Guiziou, S., Maranas, C. J., Chu, J. C. & Nemhauser, J. L. An integrase toolbox to record gene-expression during plant development. Nature communications 14, 1844, doi:10.1038/s41467-023-37607-5 (2023).

33 Chen, W., Kaur, G., Hou, L., Li, R. & Ow, D. W. Replacement of stacked transgenes in planta. Plant biotechnology journal 17, 2029–2031, doi:10.1111/pbi.13172 (2019).

34 Li, Y. et al. Recombinase-mediated gene stacking in cotton. Plant physiology 188, 1852–1865, doi:10.1093/plphys/kiac005 (2022).

35 Deich, C. et al. T7Max transcription system. Journal of Biological Engineering 17, 4, doi:10.1186/s13036-023-00323-1 (2023).

36 Franco, R. A. L. et al. Signal Amplification for Cell-Free Biosensors, an Analog-to-Digital Converter. ACS synthetic biology 12, 2819–2826, doi:10.1021/acssynbio.3c00227 (2023).

37 Chiu, W. et al. Engineered GFP as a vital reporter in plants. Current biology : CB 6, 325–330, doi:10.1016/s0960-9822(02)00483-9 (1996).

38 Müller, K. et al. A red light-controlled synthetic gene expression switch for plant systems. Molecular bioSystems 10, 1679–1688, doi:10.1039/c3mb70579j (2014).

39 Thomson, J. G., Chan, R., Thilmony, R., Yau, Y. Y. & Ow, D. W. PhiC31 recombination system demonstrates heritable germinal transmission of site-specific excision from the Arabidopsis genome. BMC biotechnology 10, 17, doi:10.1186/1472-6750-10-17 (2010).

40 Durrant, M. G. et al. Systematic discovery of recombinases for efficient integration of large DNA sequences into the human genome. Nature biotechnology 41, 488–499, doi:10.1038/s41587-022-01494-w (2023).

41 Rech, E. L., Vianna, G. R. & Aragão, F. J. High-efficiency transformation by biolistics of soybean, common bean and cotton transgenic plants. Nature protocols 3, 410–418, doi:10.1038/nprot.2008.9 (2008).

42 Wick, R. R., Judd, L. M. & Holt, K. E. Performance of neural network basecalling tools for Oxford Nanopore sequencing. Genome biology 20, 129, doi:10.1186/s13059-019-1727-y (2019).

43 Li, H. Minimap2: pairwise alignment for nucleotide sequences. *Bioinformatics (Oxford*, England) 34, 3094–3100, doi:10.1093/bioinformatics/bty191 (2018).

